# Large scale production of human blastoids amenable to modeling blastocyst development and maternal-fetal crosstalk

**DOI:** 10.1101/2022.09.14.507946

**Authors:** Leqian Yu, Toshihiko Ezashi, Yulei Wei, Jialei Duan, Deirdre Logsdon, Linfeng Zhan, Asrafun Nahar, Carlos A. Pinzon Arteaga, Lizhong Liu, Caitlen Stobbe, Mandy Katz-Jaffe, William B Schoolcraft, Lei Wang, Tao Tan, Gary C. Hon, Ye Yuan, Jun Wu

**Affiliations:** Department of Molecular Biology, University of Texas Southwestern Medical Center, Dallas, TX, USA; Colorado Center for Reproductive Medicine, Lone Tree, CO, 80124, USA; Cecil H. and Ida Green Center for Reproductive Biology Sciences, University of Texas Southwestern Medical Center, Dallas, TX, USA; State Key Laboratory of Primate Biomedical Research, Institute of Primate Translational Medicine, Kunming University of Science and Technology, Kunming, Yunnan 650500, China; Department of Obstetrics and Gynecology, University of Texas Southwestern Medical Center, Dallas, TX, USA; Department of Bioinformatics, University of Texas Southwestern Medical Center, Dallas, TX, USA; Hamon Center for Regenerative Science and Medicine, University of Texas Southwestern Medical Center, Dallas, TX, USA

**Author notes:** These authors contributed equally (Leqian Yu, Toshihiko Ezashi, Yulei Wei, Jialei Duan, Deirdre Logsdon). Correspondence (G.H.), (Y.Y.), (J.W.).

## Abstract

Recent advances in human blastoids generated from naïve pluripotent stem cells have opened a new avenue for modelling early human development and implantation. Despite the success, however, existing protocols have several limitations, e.g., the use of custom-built microwell arrays impedes wide adoption by the research community, and mass production of human blastoids is hampered by low-output or low-efficiency methods. To address these issues, here we developed an optimized protocol based on commercially available microwell plates, which enabled efficient generation of high-fidelity human blastoids at a large scale. Leveraging on the improved protocol, we identified MAPK. PI3K/AKT and mTOR signaling pathways were activated in both blastoids and blastocyst, and discovered endometrial stromal effects in promoting trophoblast cell survival, proliferation and syncytialization during extended co-culture with blastoids. Our optimized protocol will facilitate broader use of human blastoids as an accessible, perturbable, scalable, tractable, and ethical model for human blastocysts.

## INTRODUCTION

During human pre-implantation development, a blastocyst - a spherical structure made up of an outer trophectoderm (TE) epithelial layer surrounding a fluid-filled cavity that contains an inner cell mass (ICM) - forms around 5 days after fertilization. Historically, studies of pre- and peri-implantation human development have mainly relied on discarded blastocysts derived from in-vitro fertilization (IVF), which severely limit the reproducibility, scale, and types of experiments that can be performed. Human pluripotent stem cells (hPSCs) have an extended self-renewal ability and provide alternative means to study human development in a dish. Recent advances in naïve hPSC-derived blastocyst models, referred to as human blastoids, have broken new ground (Kagawa et al., 2022; Yanagida et al., 2021; Yu et al., 2021a) and provide an invaluable ethical alternative to study the early stages of human development and implantation.

By modulating several signaling pathways, we first published a two-step three-dimensional culture strategy to generate blastoids from human naïve hPSCs cultured in the 5iLA (Theunissen et al., 2014) condition (Yu et al., 2021a). Later, two independent studies reported methods to generate human blastoids from hPSCs cultured in a different naïve condition (PXGL) (Kagawa et al., 2022; Yanagida et al., 2021). Blastoids generated from both naïve conditions resembled human blastocysts in morphology, size, cell number and lineage composition, and demonstrated efficacies in studying mechanisms underlying cavity formation and first steps of human implantation (Kagawa et al., 2022; Yanagida et al., 2021; Yu et al., 2021a). Despite the similarities, there are several notable differences among protocols in terms of efficiency, scalability, accessibility, and off-target cells. Although our previous protocol is easier to be scaled-up and more accessible due to the use of commercially available AggreWell™400 microwell plates, as compared to the use of ultra-low attachment 96-well plates (Yanagida et al., 2021) and customized non-adherent hydrogel microwells (Kagawa et al., 2022), respectively, the derivation efficiency was lower and contained more off-target cells. To this end, in this study, we have developed an optimized protocol that greatly improved the derivation efficiency and fidelity of blastoids generated from 5iLA-cultured naïve hPSCs (5iLA-blastoids). The reproducibility of the optimized protocol has been validated by three independent laboratories. We also performed single cell RNA-sequencing (scRNA-Seq) for both blastoids and blastocysts using the standardized 10X Genomics Chromium platform to facilitate proper comparative analysis. Based on human blastoids generated using the optimized protocol, we studied several key signaling pathways implicated in early human development and maternal-fetal crosstalk by co-culturing blastoids with endometrial stromal cells, the cell type that intimately interact with trophoblast cells following implantation.

## RESULTS

### An optimized method to generate human blastoids from 5iLA-cultured naïve hPSCs

We previously reported the generation of 5iLA-blastoids with sequential treatments of trophoblast differentiation medium (TDM) and hypoblast differentiation medium (HDM), or vice versa (Yu et al., 2021a). We noticed that the starting cell number per microwell was a critical factor affecting blastoid formation efficiency, which is greatly influenced by cell viability following single-cell dissociation of naïve hPSCs. This varies among different PSC lines, passages, and batches, and greatly hampered the stability and efficiency of blastoid formation. To improve cell viability, we took advantage of a small-molecule cocktail termed CEPT (combination of Chroman 1, Emricasan, Polyamines, and Trans-ISRIB) that was shown to enhance the viability of hPSCs following single cell passaging (Chen et al., 2021). By supplementing HDM with the CEPT cocktail, we found starting 5iLA-hPSCs survival was greatly improved, and higher efficiency and stability in blastoid formation when compared to a ROCK inhibitor Y27632 treatment (Figures S1A and S1B), Hereinafter we designated the CEPT-supplied HDM as enhanced HDM (eHDM) (Figure 1A).

**Figure 1.**
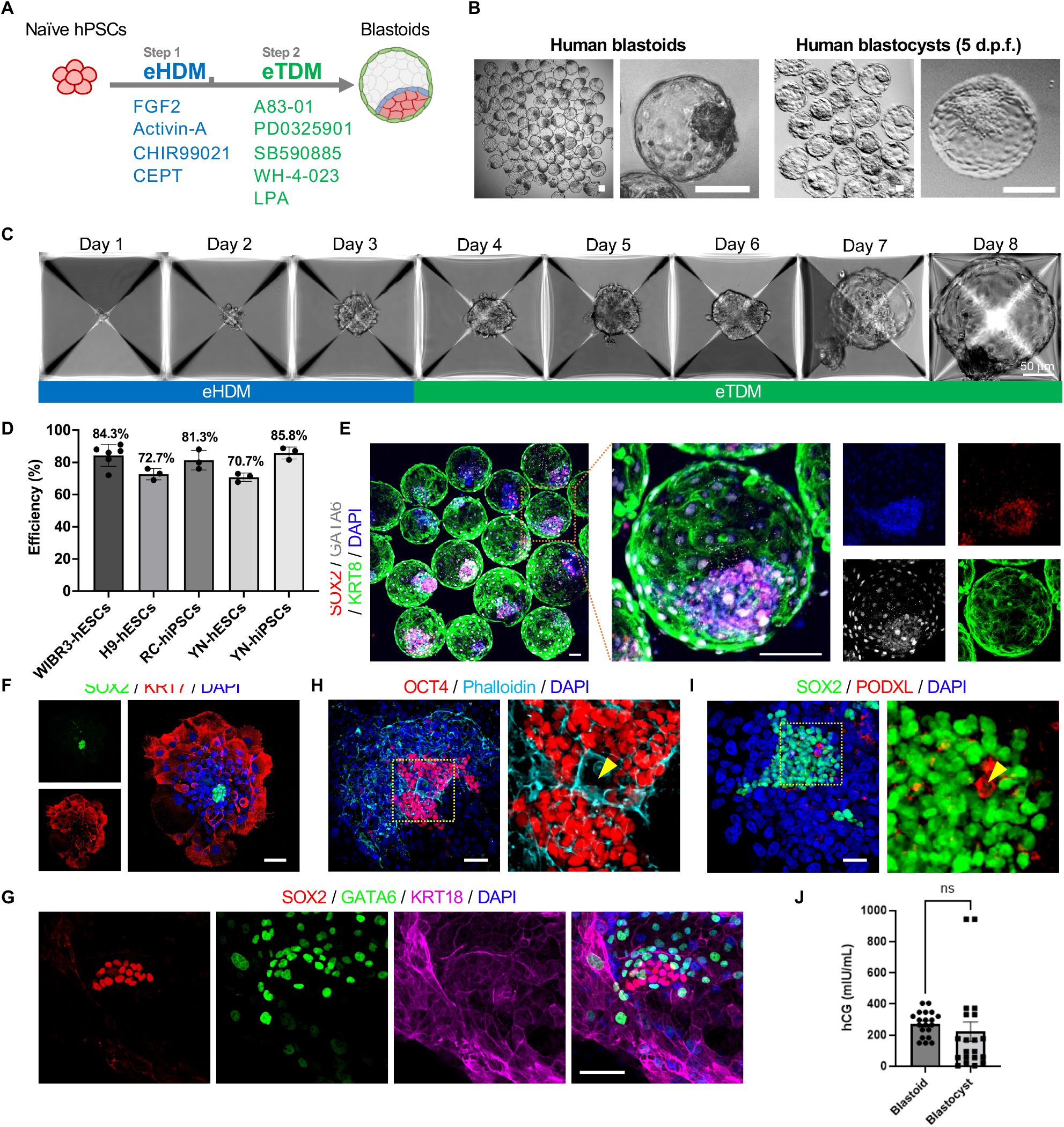
Characterization of human blastoids. (A) Schematic of the optimized blastoid formation protocol. (B) Representative phase-contrast images of human blastoids (left) and 5 d.p.f. human blastocysts (right). Scale bars, 100 μm (blastoid), 200 μm (blastocyst). (C) Representative phase-contrast images of cell aggregates at indicated time points during blastoid formation. Scale bar, 50 μm. (D) Blastoid formation efficiency from different naïve human ESC and iPSC lines. (E) Representative immunofluorescence co-staining images of SOX2, GATA6, and KRT8. Scale bar, 100 μm. (F) and (G) Representative immunofluorescence co-staining images of SOX2 and KRT7 (F), and SOX2, GATA6, and KRT18 in EBC-cultured blastoids. Scale bars, 100 μm. (H) and (I) Representative immunofluorescence co-staining images of OCT4 and phalloidin (H), and SOX2 and PODXL (I) in EBC-cultured blastoids. Higher-magnification images of the boxed areas are shown on the right. The yellow arrowheads indicate the amniotic-cavity-like structures. Scale bar, 100 μm. (J) The hCGB levels in spent media were collected from EBC-cultured human blastocysts and blastoids after 72 h in extended culture. ns, not significant.

The Hippo–YAP/TAZ signaling plays an important role in TE specification of the human blastocyst (Gerri et al., 2020), and inhibition of the Hippo pathway dramatically improved the efficiency in generating blastoid from PXGL-cultured naïve hPSCs (PXGL-blastoids)(Kagawa et al., 2022). We next asked whether 5iLA-blastoid formation efficiency could also be further improved by Hippo pathway inhibition. To this end, we added the lysophosphatidic acid (LPA), a small molecule inhibitor of the Hippo pathway, to the simplified TDM (composed of A83-01, PD0325901, SB590885 and WH-4-023), which is referred to as enhanced TDM (eTDM). After culturing starting 5iLA-hPSC aggregates (25-30 cells) for 2-3 days in eHDM followed by 5-6 days of eTDM treatment, we found the blastoid formation efficiency was dramatically increased to ~80% (Figures 1A to 1D). With this level of efficiency, we estimated more than 20,000 human blastoids could be generated in each AggreWell™400 plate, providing an ideal platform for large-scale production unparalleled by other available methods. Importantly, using the optimized protocol, human blastoids could be efficiently and robustly generated from different 5iLA-hPSC lines in three independent laboratories, demonstrating reproducibility (Figure 1D).

Together, by promoting initial cell survival and inhibiting Hippo pathway during TE induction, we have significantly improved 5iLA-blastoid formation efficiency to a comparable level reported for PXGL-blastoids (Kagawa et al., 2022).

### Characterization of human blastoids

Next, we studied the lineage composition and allocation of human blastoids generated by the optimized protocol. Immunofluorescence analyses revealed that the outer cells (Trophoblast-like cells, TLCs) stained positive for GATA3, TFAP2A, TFAP2C, TEAD4, CDX2, KRT7 and KRT8 (Figures 1E and S1C-S1H), and formed intercellular tight junctions, which are characteristic of TE cells in human blastocysts (Figure S1I). The inner cells contained ELCs (epiblast-like cells) and HLCs (hypoblast-like cells), which stained positive for epiblast (EPI) (SOX2, OCT4, and KLF17) and hypoblast (HYP) markers (GATA6), respectively, but negative for a primitive streak marker (T) (Figures 1E, S1C, S1D and S1F).

To evaluate whether human blastoids generated using the optimized protocol can form structures resembling early post-implantation human embryos following extended culture, we modified and improved an attachment culture developed for human embryos (Deglincerti et al., 2016; Ruane et al., 2022; Shahbazi et al., 2016) (Figures S2B and S2C, see Methods). After culturing human blastoids in the improved extended blastocyst culture (EBC) medium for two days and exchanging ½ volume with fresh EBC for an additional day, we observed clear segregation of EPI (SOX2 and OCT4), HYP (GATA6), and trophoblast (KRT7 and KRT18) - like lineages (Figures 1F, 1G and S2A). Notably, some blastoids self-organized into structures containing amnion-like cavities (Figures 1H and 1I). In addition, we detected human chorionic gonadotropin beta (hCGB) in the spent medium collected from EBC-cultured human blastoids, which were at comparable, if not slightly higher, levels than those from EBC-cultured human blastocysts (Figure 1J).

These results demonstrate that blastoids generated from the optimized protocol faithfully recapitulated lineage composition and allocation in human blastocysts, and retained the ability to self-organize into early post-implantation human embryo-like structures.

### Single cell transcriptomes of human blastocysts and blastoids

In our previous work, we compared single cell transcriptomes of blastoids generated using the 10X Genomics Chromium platform with published human blastocyst datasets (Yu et al., 2021a). However, at that time, all the human blastocyst single cell RNA-seq (scRNA-Seq) datasets were generated using either Smart-seq2 (Lv et al., 2019; Petropoulos et al., 2016; Xiang et al., 2020; Yanagida et al., 2021), or a custom microwell based approach (Zhou et al., 2019). Cross-platform comparison may influence the accuracy of the analysis. To minimize platform differences, in this study, we sequenced 5 day post fertilization (d.p.f.) human blastocysts (n=50) with widely accepted good quality grades above 2/3 and internal historical live birth data above 65% (Logsdon et al., 2022) side-by-side with blastoids (n=100) using the Chromium platform. To further reduce batch variation, we prepared one of the libraries on cells pooled from both human blastocysts and blastoids, and deconvoluted each cell’s origin by using sequenced genotypes (Figures S3A-S3D). We did not find pronounced batch, genotype, and gender biases for the sequenced blastocyst single cell transcriptomes (Figure S3E).

To accurately evaluate the transcriptional states of human blastoid cells, we included single cell transcriptomes from multiple publicly available datasets for comparison, including 1,529 cells from 88 human preimplantation embryos (Petropoulos et al., 2016), 555 human blastocyst cells developed up to the primitive streak anlage stage using a 3D culture condition (Xiang et al., 2020), 228 Day 5 to Day 7 human blastocyst and 267 PXGL-blastoid cells (Yanagida et al., 2021), the complete in vitro amnion dataset (Zheng et al., 2019), and 1,195 cells from a gastrulating (Carnegie stage 7, CS7) human embryo (Tyser et al., 2021) and blastoid cells from our previous work (Yu et al., 2021a) (Figures 2A and S4A-S4E). After unbiased clustering, we first visualized all human blastocyst cells from different datasets and different sequencing platforms (Figure 2, upper middle), with the cell lineage information retrieved from respective studies (Figures S4A-S4E). Single cell transcriptomes of blastocysts generated in this study (10X) clustered well with blastocyst cells from other datasets (Smart-seq2) (clusters 2, 8, 11, 12, 17, 18, 20 (Figure S4F). However, we also observed a substantial number of 10X blastocyst cells (this study) that were not represented in Smart-seq2 datasets (clusters 3, 4, 6, 7, 10, 14). These differences could be attributable to different scRNA-Seq platforms and/or transcriptional diversity among human blastocysts. Amnion-like cells were found in clusters separated from most, if not all, blastocyst and blastoid cells, which is consistent with integrated analyses from another study (Zhao et al., 2021) (Figure 2A, upper right). Based on the annotation of blastocyst cells and the expression of lineage markers, we classified blastoid cells into three lineages (Figure 2A, bottom left and middle). Notably, compared to our previous dataset, only a few cells clustered with amnion or mesoderm like cells, suggesting the improvement of blastoid fidelity using the optimized protocol (Figures 2A, bottom left and S4E). Importantly, we found the majority (~95%) of blastoid cells generated in this study clustered with blastocyst cells. Also, PXGL-blastoids (Yanagida et al., 2021) clustered well with 5iLA-blastoids, demonstrating the robustness of human blastoid models across different starting naïve hPSC conditions and different methods (Figures 2, bottom right, and S4C).

**Figure 2.**
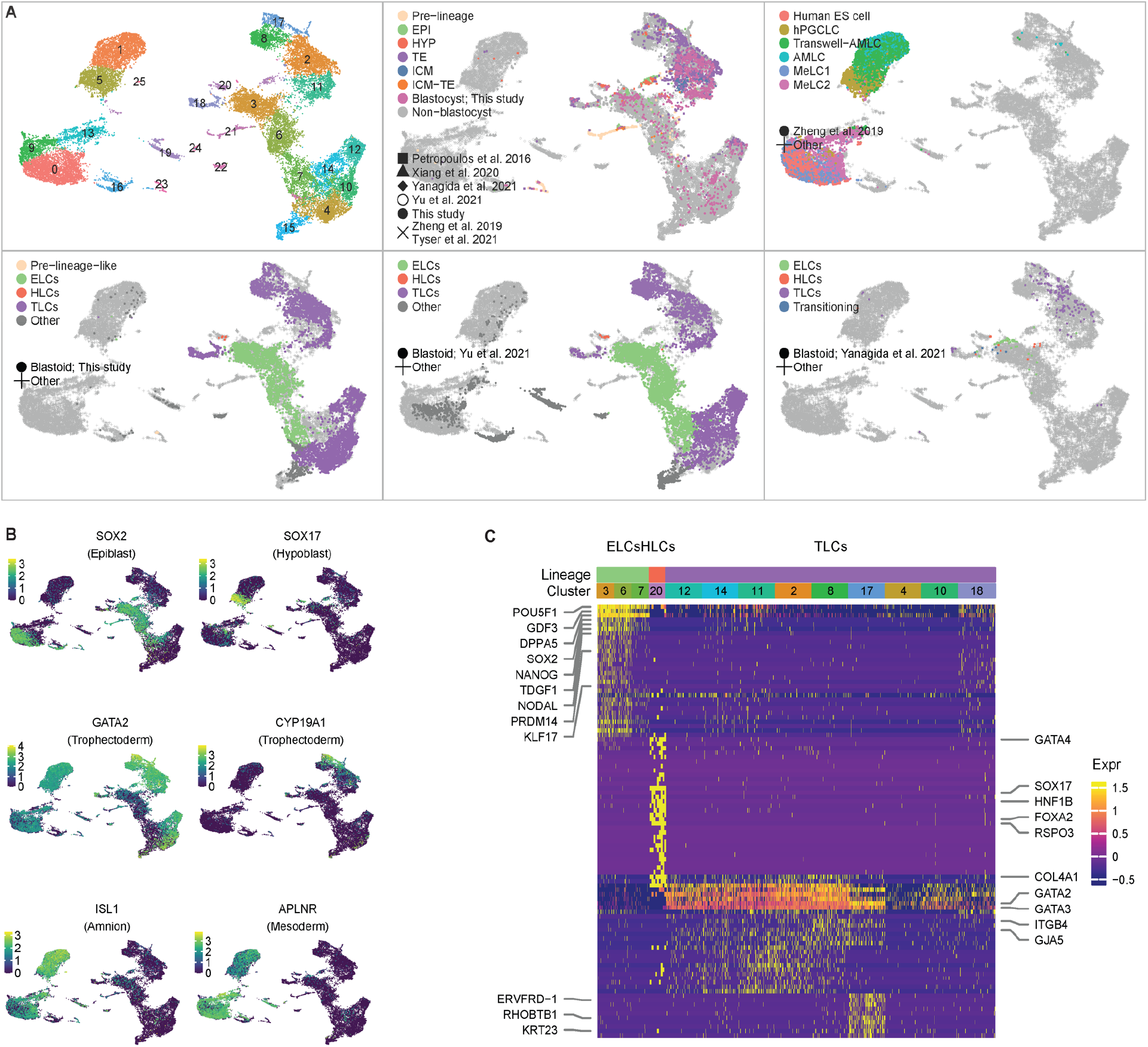
Single cell transcriptomes of human blastocysts and blastoids. (A) Joint uniform manifold approximation and projection (UMAP) embedding of single cell transcriptomes of human blastocysts, blastoids and related datasets. Upper left: colors represent different clusters. For the rest panels, shapes indicate different datasets, colors indicate cell lineages. For public datasets, cell lineage annotation is derived from their original studies. (B) Gene expression patterns of lineage markers: SOX2 (EPI), SOX17 (HYP), GATA2 (TE), CYP19A1 (polar TE), ISL1 (amnion), and APLNR (mesoderm). (C) Heatmap indicating the expression profiles of lineage-specific genes in blastoid cells. Clusters with more than 100 cells are randomly down sampled to 100 for visualization purposes.

Each of the blastoid lineages exhibited distinct gene expression profiles (Figures 2B and 2C). For instance, EPI markers SOX2, NODAL, NANOG are highly expressed in clusters 3, 6, 7; HYP markers (SOX17, GATA4, COL4A1) are highly expressed in cluster 20; and TE markers (GATA2, GATA3) are most highly expressed clusters 2, 4, 8, 10, 11, 12, 14, 17, 18. CYP19A1, a polar TE marker, is specifically expressed in cluster 17 (Figures 2B, 2C and S4G). As expected, amnion (ISL1) and mesoderm (APLNR) markers are highly expressed in non-blastoid clusters (Figure 4B).

Here, we integrated single cell reference datasets from target and other relevant cell types, as well as from different sequencing platforms, to help ensure unbiased and more accurate validation of the human blastoid model. Our analyses authenticate the human blastoids generated using the optimized protocol faithfully represent human blastocysts at the single cell transcriptome level.

### Signaling activities during human blastoid formation

Knowledge about signaling activities during early human development is limited due to restricted access to human embryos. Human blastoids provide a more accessible alternative for gaining insights into the signaling pathways involved in early human development. To this end, we collected blastoids and non-blastoid structures (aggregates without a visible cavity) and compared them to 5 d.p.f. human blastocysts using JESS, a chemiluminescent and fluorescent western blotting system. We measured the protein levels of AKT, MAPK, STAT3, P70S6, and AMPK, as well as their active (phosphorylated) forms (Figures 3A and 3B). These pathways are selected because their established roles in pluripotency and/or blastocyst development (Bell and Watson, 2013; Bessonnard et al., 2019; Bora et al., 2021; Calder et al., 2017; Hatakeyama, 2012; Murakami et al., 2004; Riley et al., 2005). Our results revealed that the ratios of p-AKT/AKT, p-MAPK/MAPK, p-P70S6/P70S6 were significantly higher in blastoids than non-blastoid structures, suggesting PI3K/AKT, MAPK and mTOR pathways play important roles during blastoid formation. In agreement, we found these signaling pathways were also activated in human blastocysts. Interestingly, the extent of signaling activation was significantly higher in human blastocysts than blastoids (Figure 3B). In contrast, we didn’t observe significant difference in the ratios of p-STAT3/STAT3 and p-AMPK/AMPK among blastoids, non-blastosid structures and blastocysts (Figures 3A and 3B).

**Figure 3.**
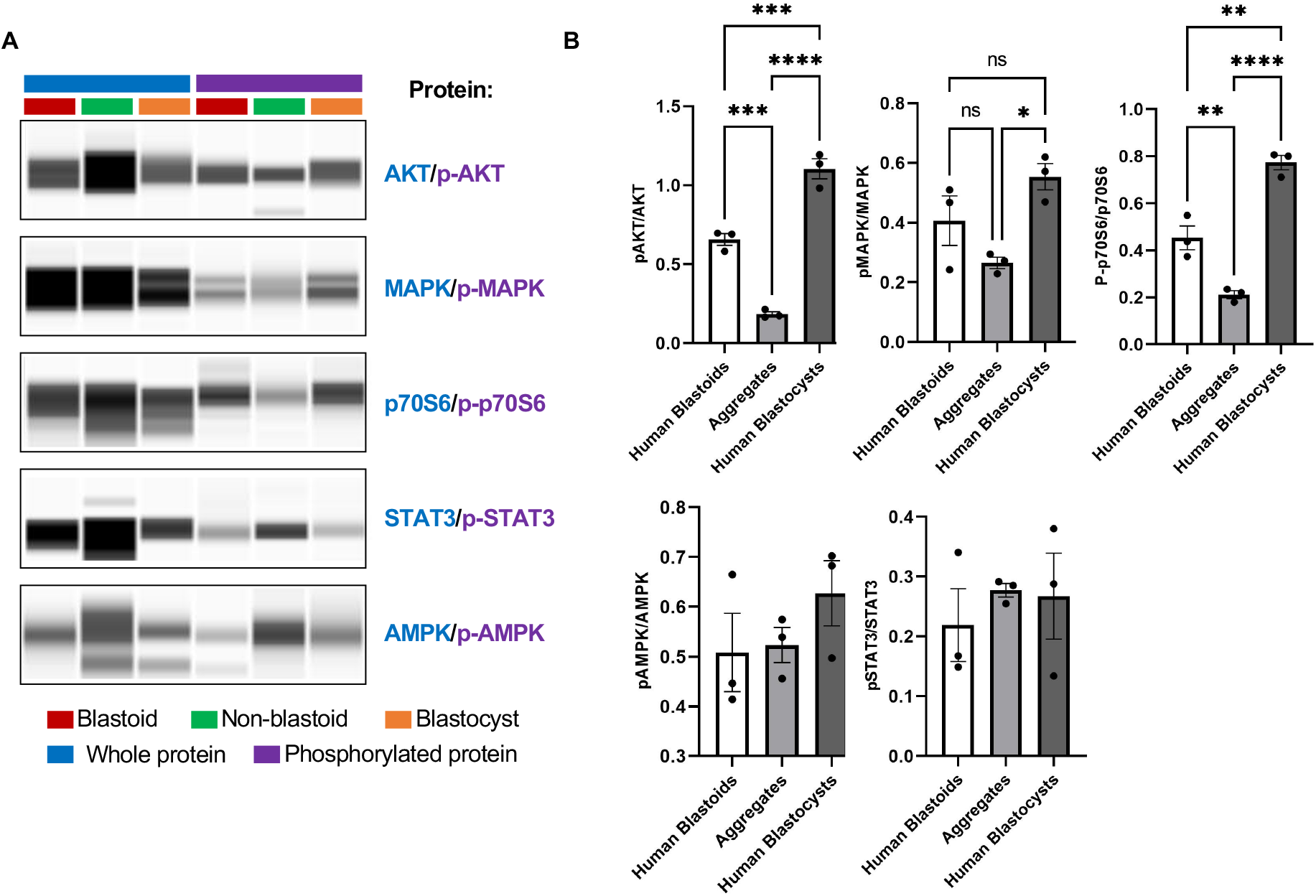
Signaling activities in human blastoids and blastocysts. (A) JESS western blotting presented as lane view for total and phosphorylated form of AKT, MAPK, p70S6, STAT3, and AMPK proteins in human blastocysts, blastoids, and non-blastoid structures. (B) Ratio of phosphorylated AKT, MAPK, p70S6, STAT3, and AMPK proteins in human blastocysts, blastoids, and non-blastoid structures. *p<0.05; **p<0.01; ***p<0.001; ****p<0.0001; ns, not significant.

### Endometrial cells promote TLCs proliferation and differentiation

During human implantation, the polar TE of a blastocyst attaches to the endometrial lining of the uterus, invades the endometrial epithelium, stroma, and the maternal circulation to form the placenta (Hustin and Schaaps, 1987). Implantation is a complex process in which several cell types are involved. Endometrial epithelium is the first maternal contact for an implanting embryo and only serves as a transient gateway for embryo implantation (Ye, 2020). Following the penetration through epithelium, the differentiation and invasion of the early human trophoblast occur in endometrial stroma in a controlled manner that ultimately result in a functional placenta (Grewal et al., 2008). A recent study established an in vitro implantation model by co-culturing human blastoids with endometrial epithelial cells (Kagawa et al., 2022). To date, however, how endometrial stromal cells influence the growth and differentiation of implanting blastoids remains elusive. To fill this gap, we established a co-culture system by growing human blastoids on immortalized primary endometrial stromal cells (IESCs) to study their crosstalk.

We first established primary cultures of endometrial stromal cells (see Methods) from patient uterine biopsies, which exhibited typical fibroblast-like morphology (Figures 4C and S6A). Because primary endometrial stromal cells have limited self-renewal ability in culture, we subjected them to immortalization by forced expression of telomerase reverse transcriptase (TERT) (Harada et al., 2003), we plated human blastoids on a hormone-stimulated monolayer of IESCs in EBC medium and compared their growth to those grown on female umbilical cord fibroblasts (herein referred to as fibroblasts) (Figure S6A) or fibronectin-coated dishes (Figures 4H and S6B). After plating, we found human blastoids quickly attached to the IESCs and readily grew outward, similar to what were observed on fibroblasts and fibronectin-coated plates (Figure 4D). After staining with the trophoblast marker GATA3, we found significantly more GATA3+ nuclei in blastoid outgrowths on both fibroblasts and IESCs than on fibronectin (Figure S6B). Interestingly, we found a significantly increased proportion of multinucleated syncytiotrophoblast (ST)-like cells (STLCs) from blastoid outgrowths on IESCs than on fibroblasts and fibronectin (Figures 4C-4E). In agreement, both hCGB fluorescent signal and the percent of hCGB+/GATA3+ nuclei were increased in blastoid outgrowth on IESCs than on fibroblasts and fibronectin (Figures 4F and 4G). When we sampled the spent medium, hCG production from blastoid outgrowths on fibronectin or IESCs was comparable and was markedly increased compared to those grown on fibroblasts (Figure S6C).

**Figure 4.**
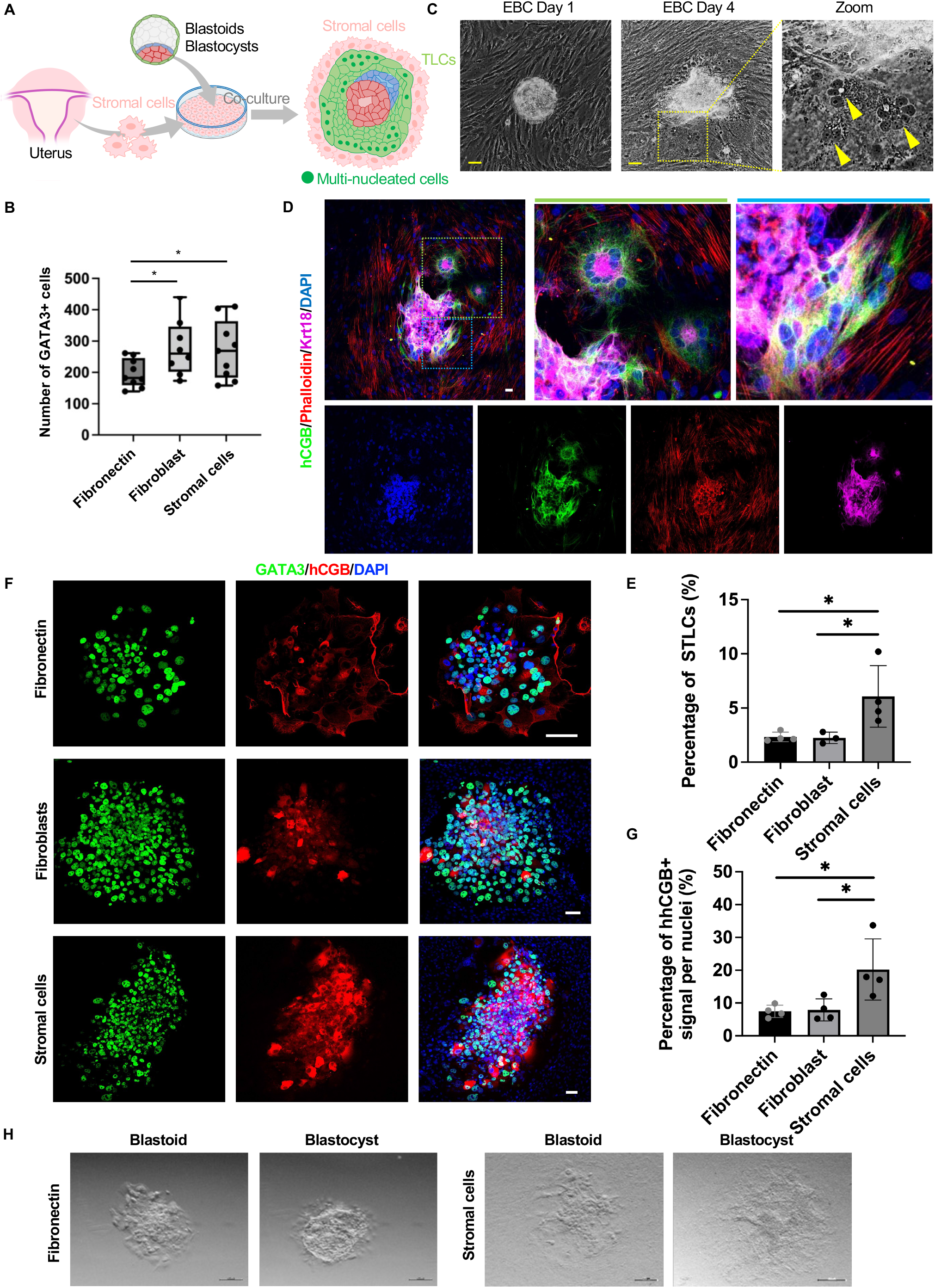
Endometrial cells promote blastoid TLCs survival and differentiation. (A) Schematic of co-culture of endometrial stromal cells and blastoids or blastocysts. (B) Cell number of GATA3+ cells in EBC-cultured blastoids on fibronectin, fibroblasts, or endometrial stromal cells after 72 h in extended culture. *p<0.05. (C) Representative phase-contrast images of days 1 and 4 EBC-cultured blastoids on endometrial stromal cells. The yellow arrowheads indicate multi-nucleated cells. Scale bars, 100 μm. (D) Representative immunofluorescence co-staining image of hCGβ, phalloidin, and KRT18 in EBC-cultured blastoids. Higher-magnification images of the boxed areas are shown on the right. Scale bar, 100 μm. (E) Percentage of the multi-nucleated STLCs of total GATA3+ cells in EBC-cultured blastoids. *p<0.05. (F) Representative immunofluorescence co-staining image of hCGβ and GAT A3 in EBC-cultured blastoids grown on different feeder conditions. Scale bar, 100 μm. (G) Percentage of the hCGB+ nuclei of total GATA3+ nuclei in EBC-cultured blastoids. *p<0.05. (H) Representative stereomicroscope images of blastoids or human blastocysts grown on fibronectin plates (left) or stromal feeder cells (right). Scale bar, 200 μm.

We then compared blastoid peri-implantation dynamics to human blastocysts. On both IESCs and fibronectin, we found most human blastocysts exhibited a 24-48 delay in attachment compared to the blastoids (Figure S6D). Late-attached blastocysts retained a large blastocoele cavity that reduced in size gradually and concurrently with trophoblast differentiation at the cell interface (Supplementary Videos 2 and 4). After 72 h, these late-attached blastocysts were largely proliferative but with reduced syncytialization and less hCGB+ cells (Figure S6F, left panel). Approximately 30% of human blastocysts were able to attach to fibronectin-coated plates or IESCs early within 24 h and exhibited reduced proliferation and early blastocoele collapse with more accelerated ST differentiation and higher number of hCGB+ cells (Figure S6F, right panel). Though trophectoderm outgrowth area was similar amongst human blastocysts and blastoids during extended culture, blastoids exhibited an attachment and outgrowth dynamic similar to the early-attached human embryos (Figures S6E and S6F, bottom panel). We also observed pronounced cell apoptosis in both peri-implantation blastoids and human blastocysts on fibronectin coated plates (Supplementary Videos 1 and 2). In sharp contrast, both blastoids and human blastocysts showed little to no apoptotic cells when grown on IESCs (Supplementary Videos 3 and 4).

Collectively, our results demonstrate endometrial stromal cells improved cell survival, encouraged proliferation, and promoted TLC syncytialization of human blastoids. Additionally, our findings reveal extended culture of human blastoids model a population of early-attached blastocysts but not the late-attached majority of human blastocyst, suggesting more room for improvements.

## DISCUSSION

Based on our previously established two-step protocol, in this study, we further optimized the efficiency, robustness, and stability of human blastoid generation from 5iLA-cultured naïve hPSCs, and demonstrated reproducibility from several human embryonic stem cell (ESC) and induced pluripotent stem cell (iPSC) lines across three independent laboratories. Two key improvements include: 1) the addition of CEPT cocktail in the HDM, which greatly enhanced the starting cell viability, and thereby ensuring the protocol’s robustness and stability; 2) the supplementation of a Hippo pathway inhibitor during trophoblast differentiation, which significantly boosted the blastoid formation efficiency and in agreement with Kagawa et al (Kagawa et al., 2022).

One of the advantages of blastoids in serving as a proxy to blastocysts for studying human peri-implantation development is that they can be produced in large quantities. Yanagida et al used non-adherent, ‘U’-bottomed 96-well plates and relied on a mouth-controlled pipette for media change, which proved to be tedious and constituted a major barrier for large-scale production (Yanagida et al., 2021). Kagawa et al, on the other hand, used a customized non-adherent hydrogel microwells (Kagawa et al., 2022), which are not readily available to the wide research community. One key strength of our protocol is the use of commercially available AggreWell™400 plates that contains 1,200 microwells in each well. Even with our previous protocol (Yu et al., 2021a) that reported lower blastoid formation efficiency (~4-29%), we could consistently generate >2,000 blastoids per AggreWell™400 plate. Here, by using the optimized method, we not only significantly improved the efficiency (~70.7%-85.8% across different cell lines) but also dramatically improved the scale-up of human blastoids generation (by estimate >20,000 blastoids can be generated per AggreWell™400 plate). Large-scale production of human blastoids opens the door for high-throughput chemical and genome-wide screens.

Another difference, and potentially an advantage of our protocol over Yanagida et al. and Kagawa et al, is that we use much fewer starting cells (25-30 cells). With additional adjustments, e.g., use of helper cells (Li et al., 2019), human blastoid can potentially be clonally generated from a single naïve hPSC. A disadvantage of our protocol is longer generation time (7-9 days) versus 3-4 days reported for PXGL-blastoids (Kagawa et al., 2022; Yanagida et al., 2021) due to the use of fewer starting cells and an additional step for directed hypoblast differentiation. For comparison, both Yanagida et al and Kagawa et al’s protocols only induced trophoblast differentiation from PXGL-hPSCs and relied on random differentiation thereafter to generate HLCs.

In both our previous and the current studies, we performed high throughput and unbiased single-cell RNA sequencing of blastoids using the standardized 10X Genomics Chromium platform. However, published scRNA-seq datasets for human blastocysts have been exclusively generated by lower throughput platforms like Smart-Seq2 that have not been commercially standardized. Problematically, the authentication of human blastoids based on cross-platform integration will introduce bias and influence the accuracy of the comparison. To address this problem, we generated a reference scRNA-Seq dataset of human blastocysts using the 10X Chromium platform. Interestingly, this dataset recovers several cell clusters that were not identified in Smart-seq2 datasets. After eliminating platform differences, blastoids generated using the optimized protocol exhibited a much higher degree of concordance (95%) with blastocysts than our previously reported blastoids (77%). In agreement with a recent study and our previous report (Zhao et al., 2021; Yu et al., 2021a), despite being a valid human blastocyst model, we found some off-target cells in our previous blastoid scRNA-Seq dataset, which were transcriptonally related to amnion- and mesoderm-like cells. These results demonstrate that, in addition to the improved efficiency, robustness, and stability, the optimized protocol also generated human blastoids with better fidelity.

Blastocyst development is accompanied by two initial cell fates determination. The outer cells are fated toward TE and the remaining cells give rise to the ICM. This is followed by ICM bifurcating into EPI and HYP. Lineage differentiation in pre-implantation embryos is orchestrated by a series of signaling pathways that are vaguely understood in humans. Among the five examined signaling pathways in this study, MAPK activation is required to regulate primitive endoderm (PE) (HYP in humans) differentiation in the ICM (Bora et al., 2021). PI3K/AKT signaling activity was detected throughout murine preimplantation development and had significant effects on the normal physiology of blastocyst (Riley et al., 2005). The phosphorylation of p70S6K is a hallmark of mTOR activity which is essential for cell growth and proliferation in early embryos (Murakami et al., 2004) and is an important mediator in maintaining PE survival and differentiation (Bessonnard et al., 2019; Bora et al., 2021). AMPK is a master regulator of cellular glucose and lipid metabolism(Hardie et al., 2012) (and its activity needs to be tightly controlled within narrow ranges to achieve optimal embryonic development to allow blastocyst formation to occur (Calder et al., 2017). JAK/STAT3 has been found to be critical for ICM formation and acquisition of naïve pluripotent state in mouse and bovine embryos (Do et al., 2013; Meng et al., 2015). It should be noted that this is the first time that these five signaling pathways were examined in human blastoids. The similarities between blastoids and human blastocysts in signaling activities demonstrate the utility of using blastoids to model human blastocyst development. Our results suggest possible roles of MAPK. PI3K/AKT and mTOR signaling pathways during human pre-implantation development, which need to be confirmed in future studies by chemical perturbation and/or genome editing of human embryos from early pre-implantation stages (Fogarty et al., 2017; Kuijk et al., 2012). The observed differences of these three pathways between blastoids and blastocysts highlight the need for future optimization to generate blastoids that can more faithfully mimic human blastocysts.

Implantation occurs when trophoblasts of the embryo attach and penetrate through the uterine epithelium and come in direct contact with the decidualized endometrial stromal cells. A rapid proliferation of the progenitor cytotrophoblast around embryonic day 8 followed by the formation of the invasive primitive ST that burrow into the endometrial stroma and secret sufficient hCG are prerequisites for establishing early pregnancy (West et al., 2019). Trophoblast proliferation, differentiation, and invasion within endometrial stroma are the early events that establish the maternal-fetal crosstalk before the formation of mature placenta. By using human blastoids as a surrogate model for human blastocysts, we demonstrated that endometrial stromal cells facilitated TLC differentiation into multi-nucleated STLCs while simultaneously promoting proliferation and mitigating apoptosis. These results are in consistent with our observations (Figures S6J-S6I) and other studies performed using human blastocysts, demonstrating the utility of human blastoids in modeling maternal-fetal crosstalk (Aberkane et al., 2018; Arjmand et al., 2016; Ruane et al., 2022). It will be interesting in future studies to use patient-specific blastoids and/or endometrial stromal cells to help understand developmental defects and early pregnancy loss.

In sum, our study showcased a robust, reproducible, and efficient method for large-scale production of high-fidelity human blastocyst-like structures from naïve hPSCs. The newly optimized protocol will facilitate broad adoption of human blastoids as an accessible, pertubable, scalable, tractable, and ethical model for human blastocysts. In this study we also identified key signaling pathways underlying blastoid formation and uncovered pro-survival and pro-differentiation effects of endometrium stromal cells on trophoblast outgrowth. These findings may help develop strategies to enhance blastoid function, thereby promoting more robust post-implantation development.

### Limitations of the study

Despite the similarities between human blastocysts and blastoids generated using the optimized protocol, there are several notable differences: 1) Human blastocysts and blastoids are generated in different culture conditions, and we found most blastoids were not able to maintain intact cavities in the presence of human blastocyst culture medium. 2) Several signaling pathways activities including PI3K/AKT, MAPK and mTOR were significantly higher in blastocysts than blastoids. 3) During extended culture in vitro, blastoids didn’t grow as robustly as most blastocysts. These differences highlight the need to further improve the blastoid model to make them more faithful surrogates for human blastocysts.

## ACKNOWLEDGMENTS

We thank R. Jaenisch and T. Theunissen for providing the naive WIBR3 (OCT4-2A-GFP) cells; We thank Ohad Gafni for providing the human primed RC-hiPSCs. We thank all the clinical embryologists in the IVF lab at Colorado Center for Reproductive Medicine (CCRM) for their contributions on human sample collection, and diagnostic lab for hCG quantification. We thank all the IVF patients who generously donated the samples for this work. J.W. is a New York Stem Cell Foundation (NYSCF)–Robertson Investigator and Virginia Murchison Linthicum Scholar in Medical Research and this work is funded by NYSCF and Discovery and Innovation Grant from the American Society for Reproductive Medicine (ASRM) Research Institute. T.T. is supported by the National Natural Science Foundation of China (82192871); Major Basic Research Project of Science and Technology of Yunnan (202001BC070001 and 202102AA100053). Y.Y. is supported by internal research funds provided by CCRM. G.H. is supported by CPRIT (RP190451), the Welch Foundation (I-1926-20170325), the Burroughs Wellcome Fund (1019804) and the Green Center for Reproductive Biology.

## AUTHOR CONTRIBUTIONS

L. Y., Y.Y., and J.W. conceived the study. L.Y. and Y.W. performed most of the human blastoid experiments at UT Southwestern, Dallas. T.E. performed human blastoid experiments and prepared immortalized primary endometrial stromal cells at CCRM, Lone Tree. L.Z. performed human blastoid experiments at Kunming University of Science and Technology, Kunming. C.S. and M.K.J. performed and organized human blastocyst cell collection for scRNA-seq at CCRM. M. K.J, and W.B.S. organized and performed endometrial tissue biopsy collection at CCRM. M.K.J, W.B.S. and Y.Y. contributed to human subject regulatory administration at CCRM. J.D. and G.H. performed scRNA-seq analyses. L.W. performed scRNA-seq library preparation. C.P.A. helped with human blastoid generation. A.N. performed the JESS experiments. D.L. performed extended culture of human blastocysts and blastoids. J.W., Y.Y., G.H. and T.T. supervised the study. L.Y. and J.W. wrote the manuscript with inputs from all authors.

---

**Figure S1.**
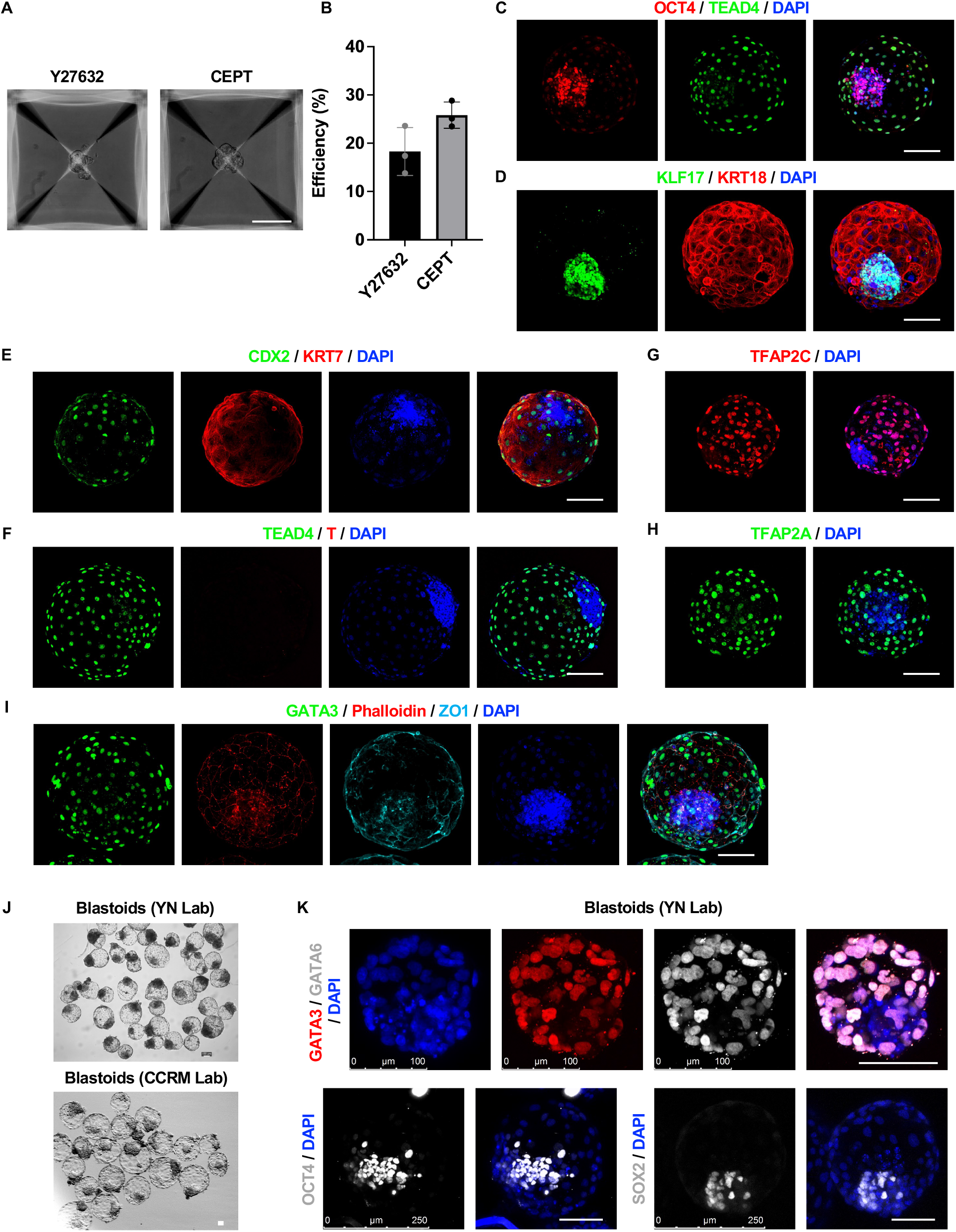
Characterization of human blastoids, relates to Figure 1. (A) Representative brightfield images of cell aggregate in microwells after 3 days of treatments with either a ROCK inhibitor Y27632 or CEPT cocktail. Scale bars, 100 μm. (B) Blastoid formation efficiency of HDM supplied with Y27632 or CEPT cocktail. (C) to (I) Representative immunofluorescence staining image of OCT4 and TEAD4 (C), KLF17 and KRT18 (D), CDX2 and KRT7 (E), TEAD4 and T (F), TCAP2C (G), TCAP2A (H) and GATA3, Phalloidin and ZO1 (I) in human blastoids. Scale bars, 100 μm. (J) Representative phase-contrast images of human blastoids generated by Lab at Yunnan, China (YN) and CCRM. Scale bars, 100 μm. (K) Representative immunofluorescence staining image of GATA3 and GATA6, SOX2, or OCT4 in human blastoids generated by YN lab. Scale bars, 100 μm.

**Figure S2.**
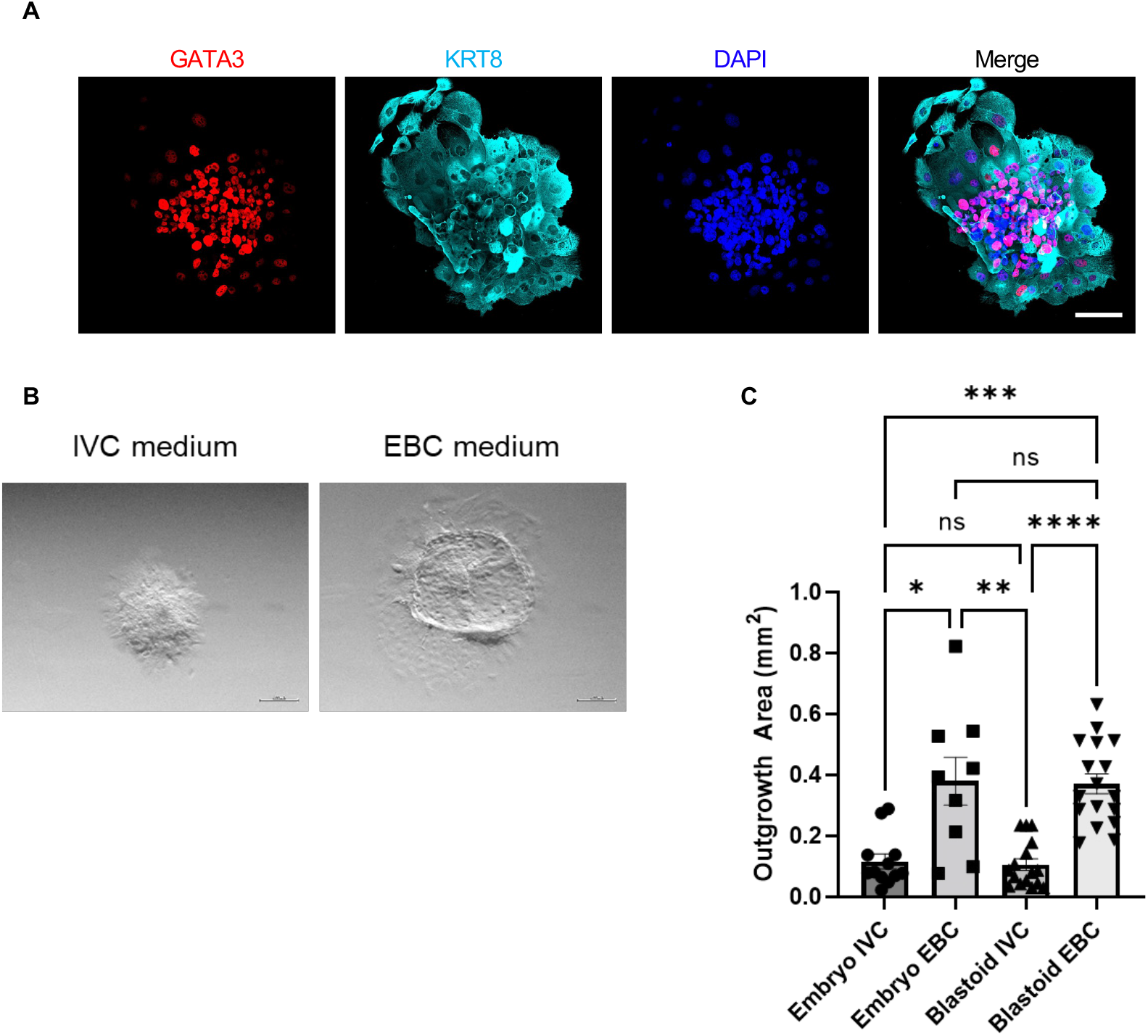
Characterization of human blastoids, relates to Figure 1. (A) Representative immunofluorescence co-staining images of GATA3 and KRT8 in EBC-cultured blastoids. Scale bar, 100 μm. (B) Representative brightfield images of extended culture of a human blastocyte in the IVC medium and EBC medium. Scale bars, 200 μm. (C) Trophoblast outgrowth area measurements of in vitro cultured human blastocysts or blastoids in the IVC medium and EBC medium after 72 h in extended culture. Kruskal-Wallis rank sum test *p<0.05; **p<0.01; ***p<0.001; ****p<0.0001; ns, not significant.

**Figure S3.**
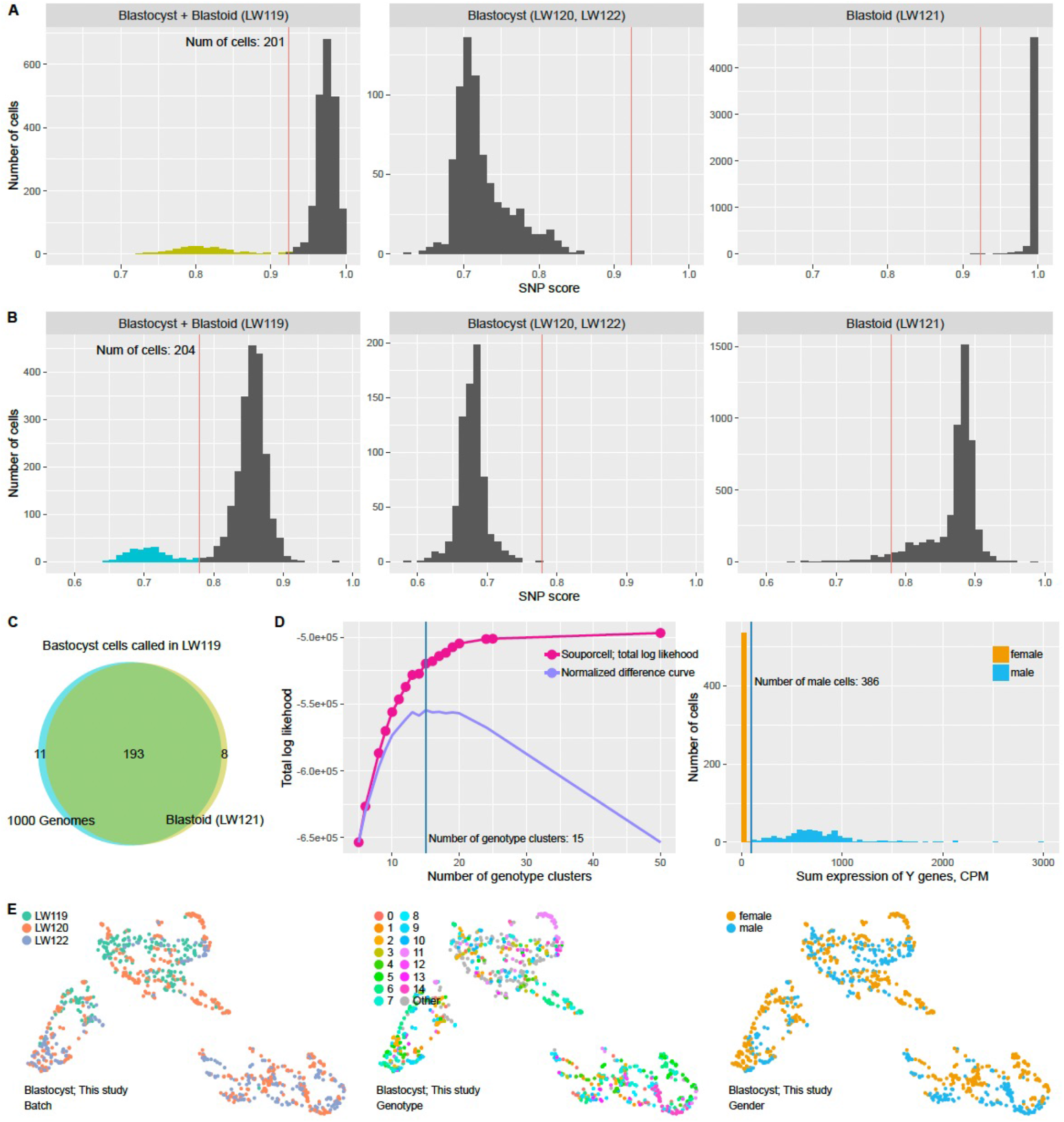
Single cell transcriptomes of human blastocysts and blastoids, relates to Figure 2. (A) (B) Distribution of SNP scores. The SNP reference is derived from de novo variant calling of blastoid cells (A) or 1000 Genomes Project (B). Thresholds were drawn on each of the plots. Blastocyst cells called in the mixed sequenced sample were colored. Separately sequenced blastocysts and blastoids were also shown. (C) High similarities of the two approaches. (D) Left, the distribution of total log likelihood of different numbers of genotype clusters. The difference curve was plotted, and the inflection point was highlighted. Right, the distribution of Y gene expressions. (E) The UMAP embedding of blastocysts generated in this study. From left to right, the colors indicate batches, genotype clusters and gender.

**Figure S4.**
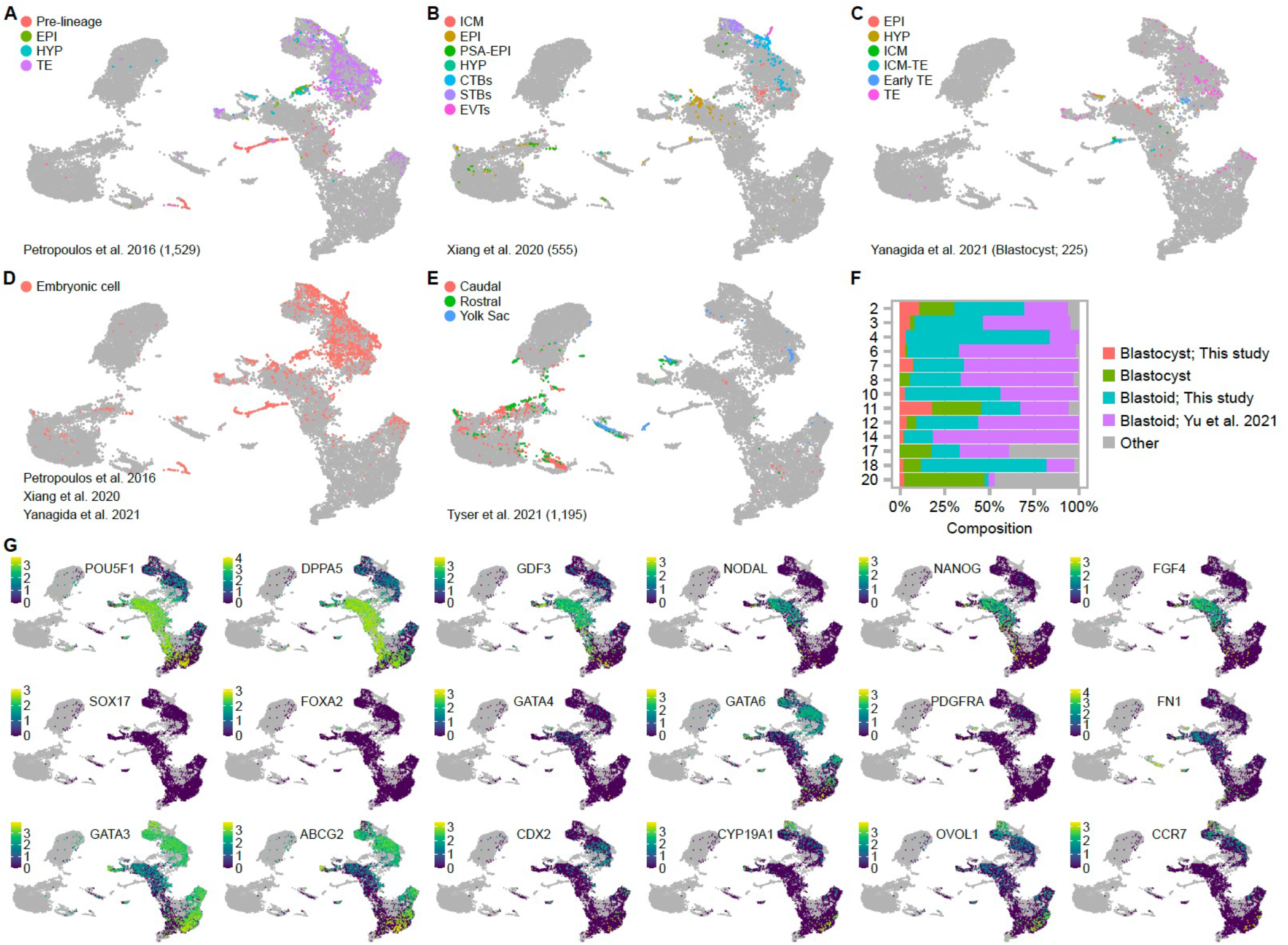
Single cell transcriptomes of human blastocysts and blastoids, relates to Figure 2. (A) - (E) The UMAP embedding of human blastoid transcriptomes. The cell lineage information is retrieved from their perspective publications. Study names are highlighted in each of the panels. Cells that are not from the studies are colored gray. (F) The cluster composition of blastoid clusters. (G) The expression patterns of EPI, HYP, and TE markers.

**Figure S5.**
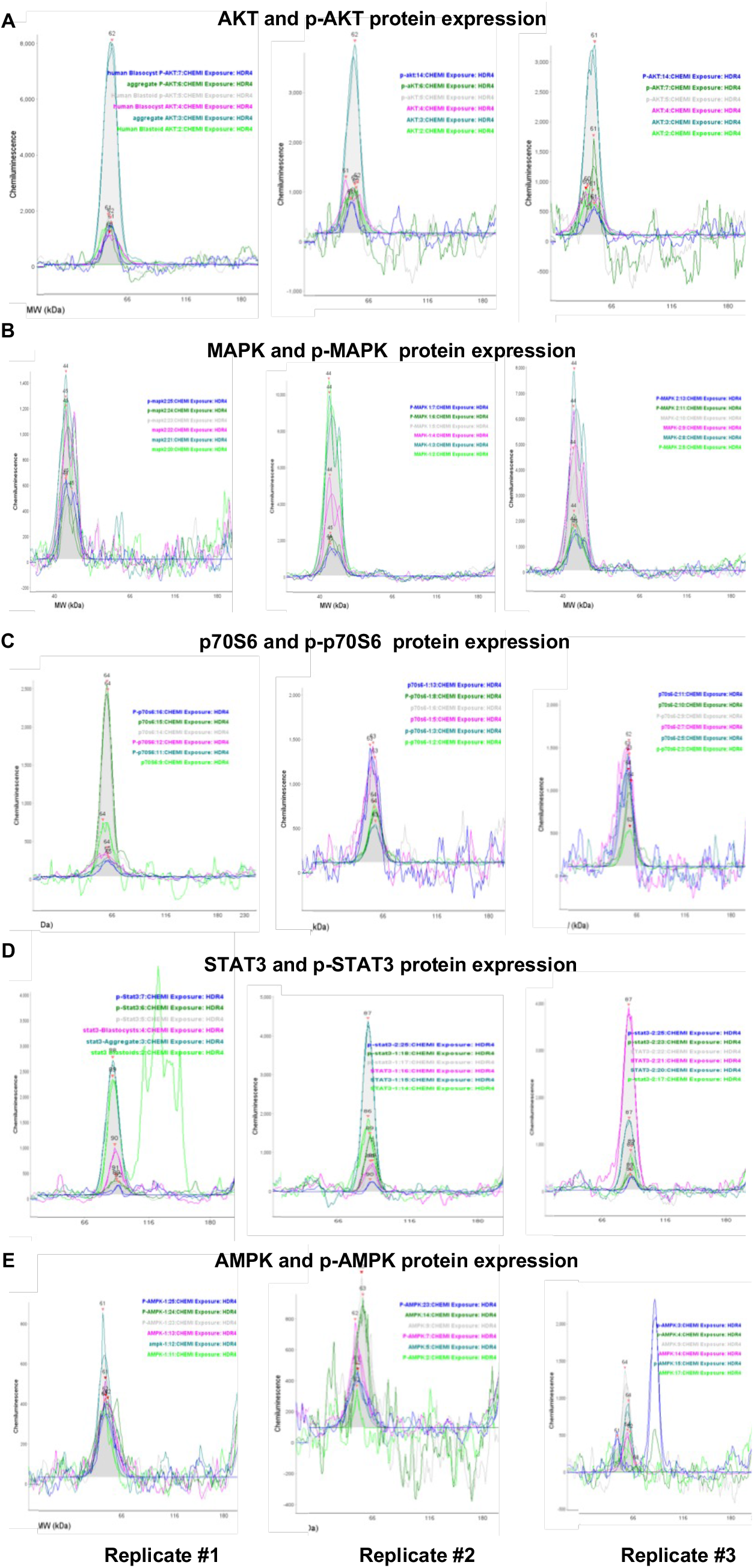
Signaling activities in human blastoids and blastocysts, relates to Figure 3. (A to E) JESS western blotting presented as electropherogram view for total and phosphorylated form of AKT, MAPK, p70S6, STAT3, and AMPK proteins in human blastocysts, blastoids, and non-blastoid structures from three independent replicates.

**Figure S6.**
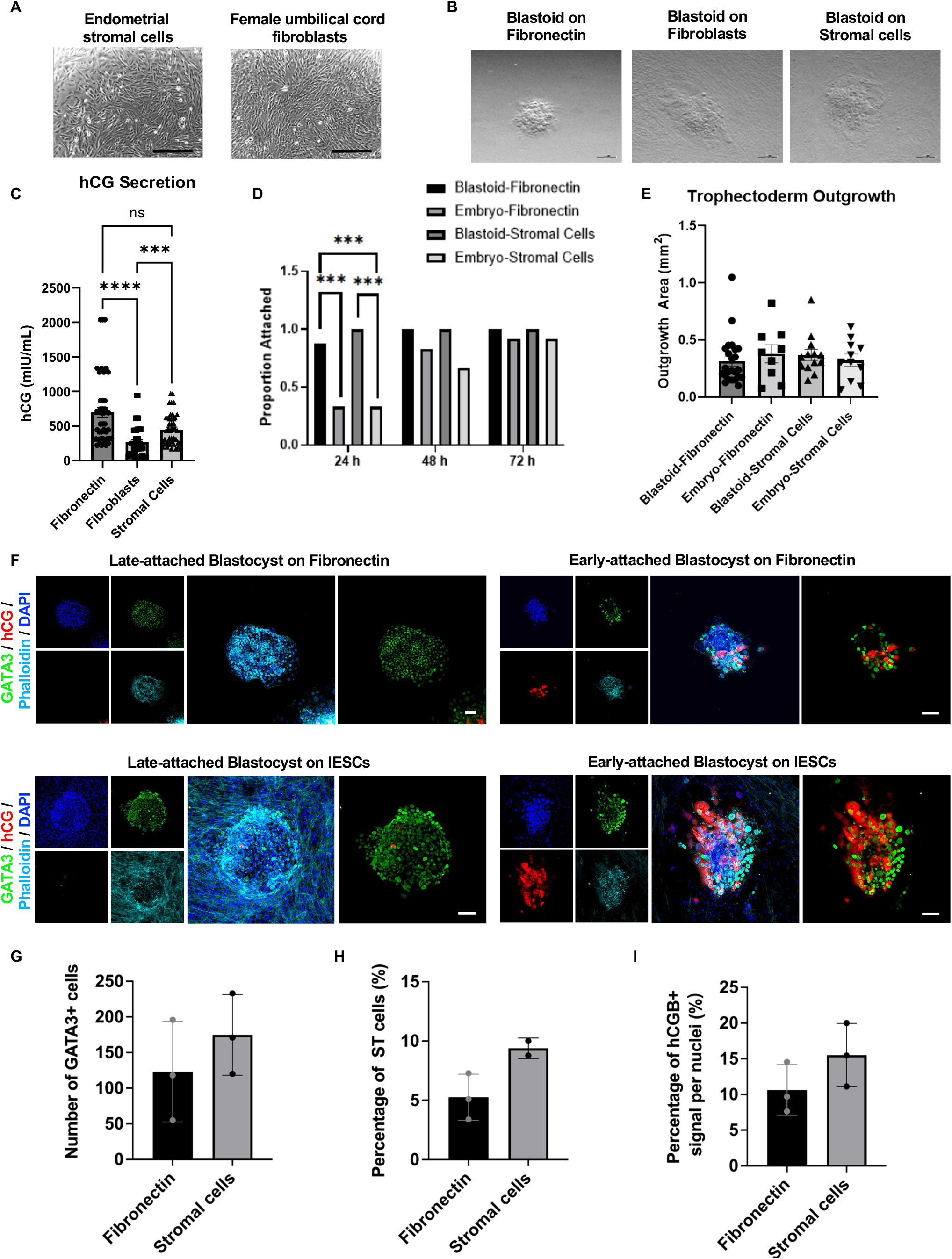
Endometrial cells promote blastoid TLCs survival and differentiation, relates to Figure 4. (A) Representative phase contrast images of endometrial stromal cells (left) and female umbilical cord fibroblasts (right). Scale bars, 200 μm. (B) Representative brightfield images of blastoids cultured on fibronectin coated plates (left), co-cultured with fibroblasts (middle), or co-cultured with endometrial stromal cells (right). Scale bars, 200 μm (C) liCGβ secretion in the spent medium (mIU/mL) of blastoids cultured on fibronectin coated plates, co-cultured with fibroblasts, or co-cultured on endometrial stromal cells. ***p<0.001; ****p<0.0001; ns, not significant. (D) Attachment rates of human blastocysts and blastoids plated on fibronectin coated plates or on endometrial stromal cells over 72 h in extended culture. ***p<0.001. (E) Trophoblast outgrowth area measurements of human blastocysts and blastoids cultured on fibronectin coated plates or co-cultured with endometrial stromal cells. (F) Representative immunofluorescence co-staining image of liCGβ and GATA3 in late-attached (left panels) or early-attached (right panels) human blastocysts cultured on fibronectin coated plates (top panels) or co-cultured with endometrial stromal cells (bottom panels). Scale bars, 100 μm. (H) Cell number of GATA3+ cells in human blastocysts (early-attached) cultured on fibronectin or co-cultured with endometrial stromal cells. (I) Percentage of multi-nucleated STs of total GATA3+ nuclei in human blastocysts (early-attached) cultured on fibronectin or co-cultured with endometrial stromal cells. (J) Ratio of the hCGB+ cells in human blastocysts (early-attached) cultured on fibronectin or co-cultured with endometrial stromal cells. Kruskal-Walllis rank sum test (C), Fisher’s Exact test (D), ***p<0.001. ****p<0.0001.

**Supplementary Table 1. Primary antibodies used in this study**

**Supplementary Video 1. Extended culture of human blastoid on fibronectin**

**Supplementary Video 2. Extended culture of human blastocyst on fibronectin**

**Supplementary Video 3. Extended culture of human blastoid on IESCs**

**Supplementary Video 4. Extended culture of human blastocyst on IESCs**

## METHODS

### Ethics statement

Most human pluripotent stem cells, endometrial cells, and blastoid experiments were performed at the UT Southwestern Medical Center and followed the 2016 Guidelines for Stem Cell Research and Clinical Translation released by the International Society for Stem Cell Research (ISSCR). Human pluripotent stem cells, endometrial cells, and blastoid experiments were reviewed and approved by the UT Southwestern Stem Cell Oversight Committee (SCRO) (registration no. 29). All human-embryo-related experiments were conducted at Colorado Center for Reproductive Medicine. The studies involving patient materials were performed following informed consent and were reviewed and approved by the WCG Institutional Review Board (WCG IRB) study no. 1179872 (blastocyst extended culture during the peri-implantation period), 20140449 (endometrial biopsies), 1332853 (blastocyst scRNA-Seq and data submission). The embryos were donated by couples undergoing in vitro fertilization (IVF). No embryos were cultured past embryonic day 12.

### Human Embryo Warming and Recovery

Vitrified day 5 or 6 human embryos were donated with informed consent following successful IVF cycles at our clinic. Embryos were warmed according to manufacturer protocols (Kitazato, Shizuoka, Japan) and left to recover in 20 μL drops on an in-house single step culture medium supplemented with 20% Quinn’s Advantage Serum Protein Substitute (SPS, Origio, Cooper Surgical, Trumbull, CT) under embryo grade culture oil (Spectrum, New Brunswick, NJ) for 2 h at 37°C and 6.5% O_2_ and 7.5% CO_2_ to mimic 5% O_2_ and 6% CO_2_ at sea level. Following a 2 h recovery, zonas were removed using a brief exposure to warmed acidic Tyrode’s solution (Sigma Aldrich, St. Louis, MO) and returned to their recovery drops before single-cell dissociation or extended culture.

### Endometrial stromal cell isolation

Endometrial cells were obtained from patient biopsy during oocyte retrieval. biopsies were washed, minced, and enzymatically digested with 0.4mg/mL Collagenase V (Sigma) and 1.25 U/mL Dispase II (Sigma). Digestion suspension was neutralized and passed through a 100 μm cell strainer to separate large glandular tissue fragments and then a 40 μm cell strainer. The filtrate from glandular fragment separation was further passed through a 10 μm cell strainer to separate stromal cells. Filtrate was centrifuged and resuspended in stromal culture 5 mL medium containing 10% fetal bovine serum (FBS, Gibco) and 1X ITS-X (ThermoFisher) in DMEM/F12 and transferred to a T25 tissue flask and incubated for 15 min. The medium suspension was then moved to a new T25 tissue flask to separate out contaminating epithelial cells.

### Endometrial stromal cell culture and plating

The isolated stromal cells were immortalized by telomerase reverse transcriptase (TERT) overexpression. Early passage (p1) of primary stroma cells were infected with Lentivirus particles for hTERT overexpression driven by EEF1A1 promoter (GeneCopoeia, LP730-025) according to the manufacture’s protocol. Cells infected with 5 MOI particles were used for the experiments within seven passages. Stromal cells were plated 3 days before extended culture for the cell expansion. On the day before extended culture, stromal cells were detached using TrypLE (ThermoFisher, Waltham, MA) and plated in ibidi dishes at 69,000 cells/cm2. Stromal cells were cultured in Advanced DMEM/F12 with 1X ITS-X (ThermoFisher), 1X GlutaMAX (ThermoFisher), and 5% fetal bovine serum (Sigma Aldrich).

### Culturing of human naïve PSCs

Naive WIBR3 human ES cells were obtained from R. Jaenisch and T. Theunissen. Primed H9-hESCs, 46XX-hESCs, RC-hiPSCs and JB-hiPSCs were cultured in mTeSR Plus medium (Stemcell technologies). Primed human PSC lines were converted to the naïve state following a previously described protocol(Theunissen et al., 2014). All naive human PSCs were cultured on mitotically inactivated mouse embryonic fibroblast (MEF) feeders at 37 °C in 5% CO_2_ and 5% O_2_. In brief, around 2 × 105 cells were plated into 6-well plates pre-coated with MEFs in 5i/L/A medium. The cells were passaged with Accumax (Stemcell technologies). The 5i/L/A medium (500 ml) was prepared using the following: 250 ml DMEM/F12 (Invitrogen), 250 ml neurobasal medium (Invitrogen), 5 ml N2 supplement (Invitrogen), 10 ml B27 supplement (Invitrogen), 1× GlutaMAX (Gibco), 1× nonessential amino acids (Gibco), 0.1 mM β-mercaptoethanol (Gibco), 0.5% penicillin–streptomycin (Gibco), 50 mg ml-1 bovine serum albumin (BSA, Sigma) and the following small molecules and cytokines: 1 μM PD0325901 (Stemgent), 0.5 μM IM-12 (Enzo), 0.5 μM SB590885 (R&D systems), 1 μM WH-4-023 (A Chemtek), 20 ng ml-1 recombinant human LIF (Peprotech), 10 ng ml-1 activin A (Peprotech) and 5 μM Y-27632 (Selleckchem).

### Optimized protocol for human blastoids generation

The protocol was optimized from our previous works(Yu et al., 2021b, 2021a). 5iLA naive human PSCs were dissociated into single cells by incubation with Accumax (Stemcell technologies) for 3 min at 37 °C. Cells were centrifugated at 200g for 5 min and collected in 5iLA medium. Then, the cell suspension was incubated in a 0.1% gelatin-coated tissue culture dish for 30 min at 37 °C in 5% CO_2_ to remove MEF feeder cells. The supernatant containing naive human PSCs was collected and passed through a 40-μm cell strainer, and cells were counted. Meanwhile, the AggreWell-400 (STEMCELL Technologies) was prepared according to the manufacturer’s instructions. In brief, wells were rinsed with anti-adhesion rinsing solution (STEMCELL Technologies), centrifuged at max speed for 5 min, and then incubated at room temperature for 10 min. After incubation, wells were washed with eHDM once and 0.5 ml fresh eHDM was added. The corresponding number of cells (25 cells / microwell for WIBR3-hESCs, H9-hESCs. And YN-hESCs, 30 cells / microwell for RC-hiPSCs and YN-hiPSCs, and for other cell lines the starting number needs to be optimized) were resuspended in 1 ml eHDM and seeded into one well of a prepared AggreWell-400 24-well plate. The plate was centrifuged at 200g for 1 min and cultured at 37 °C in 5% CO_2_ and 5% O_2_, the day of cell plating was designated as day 0. The medium was half changed (remove 1 ml old medium were removed and 1ml fresh medium were added) with eHDM on day 1 and day 2. On day 3, carefully remove as much eHDM as possible, and then add 1.5 ml of eTDM. Repeat this step one additional time to help completely remove the remaining eHDM. On the remaining days, the fresh eTDM were half changed every two days. The human blastoids usually formed after around five to six days of culture in eTDM. All blastoids were manually isolated using a mouth pipette under a stereomicroscope for downstream experiments.

The eHDM was prepared using the following: 1:1 (v/v) mixture of DMEM/F12 and neurobasal medium, 1× N2 supplement, 1× B27 supplement, 1 × GlutaMAX, 1× nonessential amino acids, 0.1 mM β-mercaptoethanol, 0.5% penicillin–streptomycin, 20 ng ml-1 bFGF (Peprotech), 20 ng ml-1 activin A, 3 μM CHIR99021 and CEPT cocktail [50 nM Chroman 1 (MedChem Express), 5 μM Emricasan (Selleckchem), 1X polyamine supplement (Sigma), and 0.7 μM TransISRIB (Tocris)].

The eTDM was prepared using the following: 1:1 (v/v) mixture of DMEM/F12 and neurobasal medium, 0.5× N2 supplement, 0.5× B27 supplement, 0.5% ITS-X, 0.5× GlutaMAX, 0.5× nonessential amino acids, 0.1 mM β-mercaptoethanol, 0.5% knockout serum replacement (KSR, Gibco), 0.5% penicillin–streptomycin, 2 μM PD0325901, 1 μM A83-01, 0.5 μM SB590885, 1 μM WH-4-023, 10 ng ml-1 recombinant human LIF, and 0.5 μM LPA.

### Human blastocyst and blastoid extended culture

Embryos and blastocysts were placed into 300 μL of extended blastocyst culture (EBC) medium containing DMEM/F12 and Neurobasal (Thermo Fisher, 21103049) with N2 supplement (Thermo Fisher, 17502048), B27 supplement (ThermoFisher, 17504044), GlutaMAX (Gibco, 35050061), non-essential amino acids (ThermoFisher, 11140050), 0.1 mM β -mercaptoethanol, 15% fetal bovine serum (Hyclone, SH30070.02), 8 nM water-soluble estradiol (E4389), 200 ng/mL progesterone (P7556), 1 mM sodium pyruvate (SigmaP4562), 5 mg/mL Gentamicin (Cellgro, 30-005-CR), 50 nM Chroman 1 (MedChem Express, HY-15392), 5 μM Emricasan (Selleckchem, S7775), 1X polyamine supplement (P8483), and 0.7 μM TransISRIB (Tocris, 5284). All extended culture experiments were completed in 8-well ibi-Treat dishes (Ibidi, Martinsried, Germany) coated in fibronectin from human plasma (F0895). Embryos and blastoids were cultured for 72 h in the extended culture system on either fibronectin-coated ibidi dishes or ibidi dishes seeded with endometrial stromal cells. After 48 h, ½ medium volume was exchanged with fresh EBC. After 72 h, samples were fixed with 4% paraformaldehyde (Electron Microscopy Sciences, 30525-89-4).

### Human Chorionic Gonadotropin Quantification

150 μL of spent medium was collected and snap frozen in LN2 upon the conclusion of culture and stored in −80°C. Samples were analyzed for human chorionic gonadotropin (hCG) using electrochemiluminescence on a Roche Cobas e601 analyzer and is reported in mIU/mL.

### Outgrowth Area Measurement

Images of peri-implantation blastoids and embryos were captured using an Olympus DP22 (Olympus, Melville, NY) camera fixed to a stereomicroscope. In ImageJ, trophectoderm outgrowth was measured with the freehand selection tool and areas were calculated using the scale bar as a reference.

### Jess Western Blotting

Human blastoids with cavity, stem cell aggregates that did not form a cavity from the same batch, and human blastocysts with similar morphology with blastoids were flash frozen in groups of 10 with minimal amount of DPBS-0.1% PVP per tube using liquid nitrogen. Samples were stored at −80°C until used for protein expression analysis. On the day of running Jess western blotting, samples were thawed on ice and 100 μl of RIPA buffer (Sigma-Aldrich, MO, USA) supplemented with protease and phosphatase inhibitor (Sigma-Aldrich, MO, USA) was added to lyse the samples. Samples were then subjected to ultrasonication followed by centrifugation at 14,000 × g for 10 minutes to remove cellular debris. Supernatants were collected from the lysates and quantified by a Bio-Rad DC kit assay, and 500 ng of the protein was used for Jess western blotting analysis following the Manufacturer’s Instructions (ProteinSimple, San Jose, CA). The experiments were performed in triplicate and data was analyzed by one-way ANOVA with post-hoc Tukey’s multiple comparison test. Primary antibody information is included in Supplementary Table 1.

### Immunofluorescence Staining

Human blastocysts and blastoids were fixed with 4% paraformaldehyde (PFA) in PBS for 20 min at room temperature and permeabilized with 0.5% Triton X-100 in PBS for 1 h. Samples were then blocked with blocking buffer (PBS containing 5% FBS, 4% BSA and 0.1% Tween-20) at room temperature for 1 h, or overnight at 4 °C. Primary antibodies diluted in blocking buffer were applied to samples and incubated overnight at 4 °C. Samples were washed three times with PBS containing 0.1% Tween-20 (PBS-T), followed by incubation with fluorescently conjugated secondary antibodies diluted in blocking buffer for overnight at 4 °C. Samples were washed three times with PBS-T. Finally, cells were counterstained with 300 nM 4’,6-diamidino-2-phenylindole (DAPI) solution at room temperature for 10 mins. Phalloidin was directly stained along with other secondary antibodies in the blocking buffer. Samples were imaged using a fluorescence (Revolve, Echo) or a confocal microscope (A1R, Nikon). The cell numbers were counted manually using the confocal microscope (A1R, Nikon). Primary antibody information is included in Supplementary Table 1.

### scRNA-seq library preparation

Fifty human blastocysts and 100 blastoids with similar morphologies were selected for sequencing. Human blastocysts and blastoids were manually pooled using a mouth pipette, washed three times in PBS and then dissociated with trypsin-EDTA (0.25%) (Gibco) at 37 °C in 5% CO_2_ for 40 min followed by agitation using a mouth pipette under a stereoscope. Cell number and viability were examined via trypan blue assay on an automatic cell counter (product information needed). Single cell suspensions (500-1000 cells/mL) were loaded into a 10x Genomics Chromium Chip following the manufacturer’s instruction 1. (10x Genomics, Pleasanton, CA, Chromium Next GEM Single Cell 3□ GEM, Library & Gel Bead Kit v3.1, catalog number 1000121).

### Sequencing, base calling and fastq demultiplexing

Single cell libraries were sequenced using Illumina NextSeq 500/550 and NovaSeq 6000 sequencing systems (Illumina, San Diego, CA). The generated binary base call (BCL) files were demultiplexed and converted to standard FASTQ files using the mkfastq function from 10x Genomics’ Cell Ranger pipeline (version 6.1.2).

### Preprocessing single-cell data

Fastq files were mapped to the human GRCh38 reference (version 3.0.0, downloaded from 10x website) using 10x Genomics’ Cell Ranger pipeline (version 6.1.2) with default parameters. For blastoid samples, the number of cells expected was set to 10,000. For human blastocyst samples, cell-associated barcodes were determined based on the inflection point of the UMI saturation curves. Scrublet was used to detect multiplets for blastoid samples (version 0.2.1). Cells with more than 20% of UMIs from mitochondrial genomes and less than 200 genes detected were discarded.

Raw fastq files of public datasets were downloaded and re-processed to minimize platform and processing differences as described in Yu et al.(Yu et al., 2021a). The cell lineage annotations of the public datasets were derived from their perspective publications (see https://github.com/jlduan/replica for details).

### Genotyping

To separate the mix-sequenced blastocyst and blastoid cells, two approaches were used. First, the genotypic variant calling of the 3,202 samples from the 1000 Genomes Project was downloaded and only SNPs with more than 10% allelic frequency on chromosomes 1 to 22 and X were kept(Byrska-Bishop et al., 2021). This SNP list served as the reference genotype. Individual cells from separately sequenced blastoid samples were compared against the reference genotype, and a consensus genotype of the blastoid cells was summarized. Then the cells from the mix-sequenced sample were compared against the blastoid consensus genotype and scored based on the percentage of SNPs that agree with it. For the second approach, instead of using the genotypes from the 1000 Genomes Project, the reference genotype of blastoid cells was called using HaplotypeCaller module from GATK (version 4.2.4.1) and homogeneous SNPs with more than 3 reads support were kept as blastoid consensus genotypes(Poplin et al., 2018). Prior to the variant calling, reads contributing to real cells from the alignment file generated by 10x Genomics’ Cell Ranger were processed with GATK’s SplitNCigarReads module with default parameters.

The distribution of genotype scores was plotted, and the cutoff was determined using Gaussian mixture distribution model from scikit-learn (version 1.0.2) with likelihood set to 0.9.

Single cell genotype clustering was performed using souporcell (version 2.4) with a range of different numbers of clusters. And the total log likelihood of each run was recorded, and an elbow plot was generated(Heaton et al., 2020). The number of major genotypes was determined based on the inflection point of the elbow curve.

### Dimensionality reduction

Reprocessed single cell data from selected public and our previous studies were combined with newly generated data(Kagawa et al., 2022; Petropoulos et al., 2016; Tyser et al., 2021; Xiang et al., 2020; Yanagida et al., 2021; Yu et al., 2021a). The combined expression matrix was median-normalized, natural-log-transformed with the addition of a pseudocount of 1 and standardized per gene prior to principal component analysis (PCA). sc.tl.pca from Scanpy package (version 1.9.1) was used to compute the first 100 components (Wolf et al., 2018).

### Batch correction

The python implementation harmonypy (version 0.0.5, https://github.com/slowkow/harmonypy) of Harmony algorithm was used to remove the batch effect(Korsunsky et al., 2019). Each 10x Genomics run was considered as one batch, Smart-Seq2 libraries from individual studies were treated as individual batches.

### Uniform Manifold Approximation and Projection (UMAP)

Batch corrected principal components were used as input for UMAP (version 0.5.3) with default parameters(McInnes et al., 2018).

### Clustering

Single cells were clustered using the leiden algorithm implemented in the Scanpy package (version 1.9.1)(Wolf et al., 2018). Briefly, the batch corrected principal components were used as input for sc.pp.neighbors function from Scanpy package to compute a neighborhood graph of observations (n_neighbors=30). Then, leiden algorithm was used to cluster single cells with default parameters.

## Code availability

The code used in this project is provided at https://github.com/jlduan/blastoidverse.

## Data availability

Single cell RNA-seq data generated in this study have been deposited in the Gene Expression Omnibus (GEO) with accession number GSE210962.

